# Patterns of genetic differentiation and the footprints of historical migrations in the Iberian Peninsula

**DOI:** 10.1101/250191

**Authors:** Clare Bycroft, Ceres Fernandez-Rozadilla, Clara Ruiz-Ponte, Inés Quintela-García, Ángel Carracedo, Peter Donnelly, Simon Myers

**Author notes:** These authors jointly directed this work (Peter Donnelly, Simon Myers).

## Abstract

Genetic differences within or between human populations (population structure) has been studied using a variety of approaches over many years. Recently there has been an increasing focus on studying genetic differentiation at fine geographic scales, such as within countries. Identifying such structure allows the study of recent population history, and identifies the potential for confounding in association studies, particularly when testing rare, often recently arisen variants. The Iberian Peninsula is linguistically diverse, has a complex demographic history, and is unique among European regions in having a centuries-long period of Muslim rule. Previous genetic studies of Spain have examined either a small fraction of the genome or only a few Spanish regions. Thus, the overall pattern of fine-scale population structure within Spain remains uncharacterised. Here we analyse genome-wide genotyping array data for 1,413 Spanish individuals sampled from all regions of Spain. We identify extensive fine-scale structure, down to unprecedented scales, smaller than 10 Km in some places. We observe a major axis of genetic differentiation that runs from east to west of the peninsula. In contrast, we observe remarkable genetic similarity in the north-south direction, and evidence of historical north-south population movement. Finally, without making particular prior assumptions about source populations, we show that modern Spanish people have regionally varying fractions of ancestry from a group most similar to modern north Moroccans. The north African ancestry results from an admixture event, which we date to 860 - 1120 CE, corresponding to the early half of Muslim rule. Our results indicate that it is possible to discern clear genetic impacts of the Muslim conquest and population movements associated with the subsequent Reconquista.

## 1 Introduction

Genetic differences within or between human populations (population structure) has been studied using a variety of approaches over many years^1-5^. Recently there has been an increasing focus on studying genetic differentiation at fine geographic scales, such as within countries^6-8^. Identifying such structure allows the study of recent population history, and identifies the potential for confounding in association studies, particularly when testing rare, often recently arisen variants^9^. The Iberian Peninsula is linguistically diverse, has a complex demographic history, and is unique among European regions in having a centuries-long period of Muslim rule^10^. Previous genetic studies of Spain have examined either a small fraction of the genome^12-14^ or only a few Spanish regions^15,16^. Thus, the overall pattern of fine-scale population structure within Spain remains uncharacterised. Here we analyse genome-wide genotyping array data for 1,413 Spanish individuals sampled from all regions of Spain. We identify extensive fine-scale structure, down to unprecedented scales, smaller than 10 Km in some places. We observe a major axis of genetic differentiation that runs from east to west of the peninsula. In contrast, we observe remarkable genetic similarity in the north-south direction, and evidence of historical north-south population movement. Finally, without making particular prior assumptions about source populations, we show that modern Spanish people have regionally varying fractions of ancestry from a group most similar to modern north Moroccans. The north African ancestry results from an admixture event, which we date to 860 - 1120 CE, corresponding to the early half of Muslim rule. Our results indicate that it is possible to discern clear genetic impacts of the Muslim conquest and population movements associated with the subsequent *Reconquista.*

## 2 Results

We analysed phased genotyping array data for 1,413 Spanish individuals typed at 693,092 autosomal single nucleotide polymorphisms (SNPs) after quality control (Methods). We applied fineSTRUCTURE^17^ to these data to infer clusters of individuals with similar patterns of shared ancestry (Methods). fineSTRUCTURE inferred 145 distinct clusters, along with a hierarchical tree describing relationships between the clusters (**Figure 1a**; Methods). We used genetic data only in the inference, but explored the relationship between genetic structure and geography using a subset of 726 individuals for whom geographic information was available and all four grandparents were born within 80 Km of the centroid of their birthplaces. **Figure 1b** represents each of these individuals as a point on a map of Spain, located at the centroid of their grandparents’ birthplaces and labelled according to their cluster assignment after combining small clusters at the bottom of the tree (Methods). Their grandparents were likely to have been born in the early 1900s (median birth-year of the cohort is 1941), so the spatial distribution of genetic structure described in this study would reflect that of Spain around that time.

**Figure 1.**
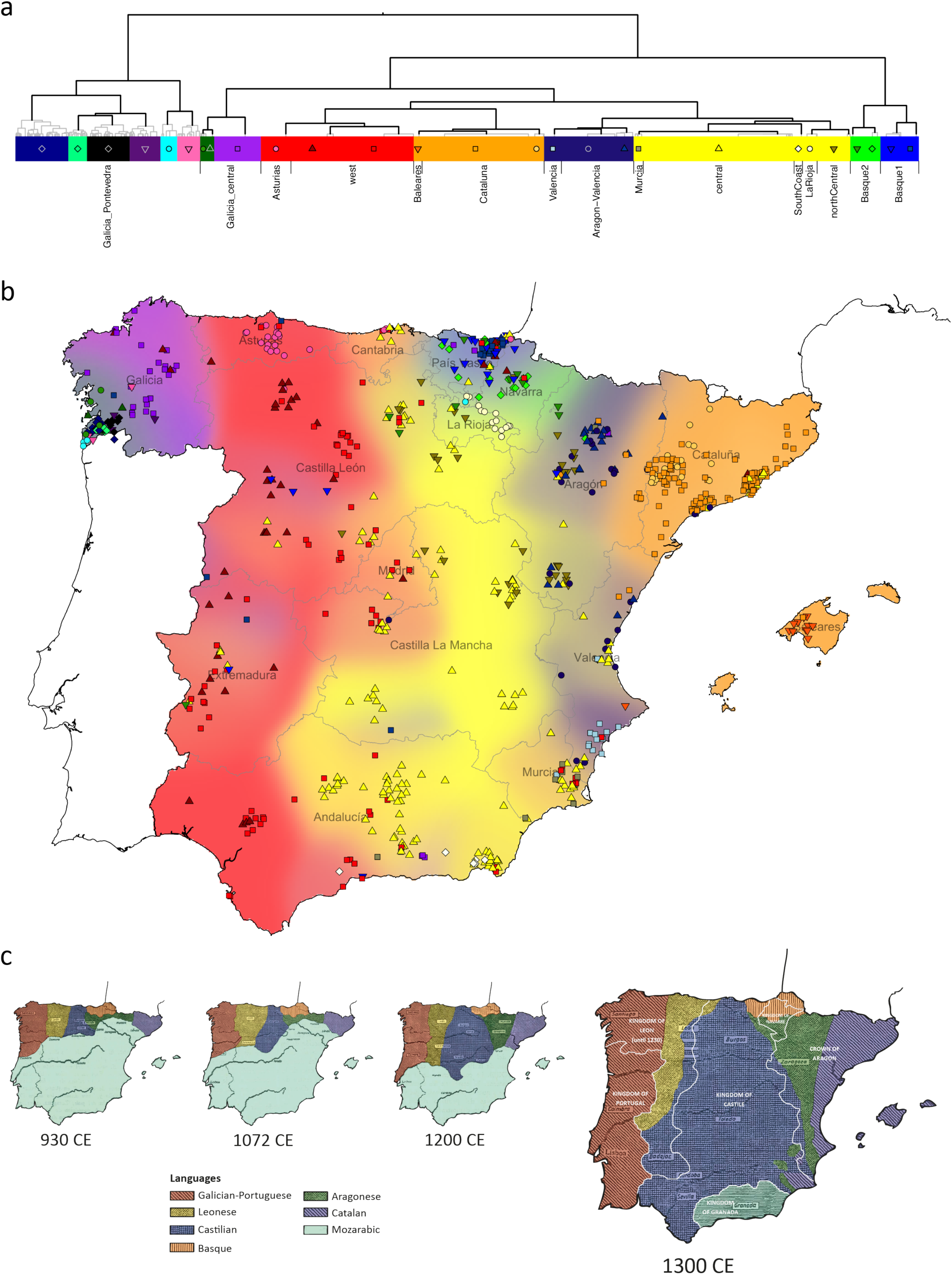
Spanish individuals grouped into clusters using genetic data only. **(a)** Binary tree showing the inferred hierarchical relationships between clusters. The colours and points correspond to each cluster as shown on the map, and the length of the coloured rectangles is proportional to the number of individuals assigned to that cluster. We combined some small clusters (Methods) and the thick black branches indicate the clades of the tree that we visualise in the map. We have labeled clusters according to the approximate location of most of their members, but geographic data was not used in the inference. **(b)** Each individual is represented by a point placed at (or close to) the centroid of their grandparents’ birthplaces. On this map we only show the individuals for whom all four grandparents were born within 80km of their average birthplace, although the data for all individuals were used in the fineSTRUCTURE inference. The background is coloured according to the spatial densities of each cluster at the level of the tree where there are 14 clusters (see Methods). The colour and symbol of each point corresponds to the cluster the individual was assigned to at a lower level of the tree, as shown in (a). The labels and boundaries of Spain’s Autonomous Communities are also shown. **(c)** A representation of changes in the linguistic and political boundaries in Iberia from ∼930 to 1300 CE, adapted from maps by Baldinger^11^. The first three maps show changes in linguistic regions only, and the last map includes political boundaries. Different linguistic areas are shown with the colours and shading, and political boundaries with white borders. Only the colours and labels of the Christian kingdoms have been added to aid visualisation.

These results reveal patterns of rich fine-scale population structure in Spain. At the coarsest level of genetic differentiation (i.e. two clusters at the top of the hierarchy) individuals located in a small region in south-west Galicia are separated from those in the rest of Spain. The next level separates individuals located primarily in the Basque regions in the north (País Vasco and Navarra) from the rest of Spain. Further down the tree (background colours in **Figure 1b**) many of the clusters closely follow the east-west boundaries of Spain’s Autonomous Communities, especially in the north of Spain. However, in the north-south direction several clusters cross boundaries of multiple Autonomous Communities. Overall, the major axis of genetic differentiation runs from east to west, while conversely there is remarkable genetic similarity on the north-south direction. In a complementary analysis that included Portugal, although fewer SNPs (Methods), Portuguese individuals co-clustered with individuals in Galicia (**Supplementary Figure 1a**), showing that this pattern extends across the whole Iberian Peninsula. Indeed, rather than solely reflecting modern-day political boundaries, the broad-scale genetic structure of the region is strikingly similar to the linguistic frontiers present in the Iberian Peninsula around 1300 CE (**Figure 1c**).

Although some geographically dispersed clusters (e.g. ‘central’ and ‘west’) remain largely intact at the bottom of the hierarchical tree (**Supplementary Figure 1b**) many of the clusters that emerge further down tree involve greater geographical localisation. By far the strongest sub-structure is seen within a single province in Galicia, Pontevedra, which contains almost half of the inferred clusters in all of Spain (**Figure 1a**). This ‘ultra-fine’ structure is seen across scales of less than 10 Km and the clusters align with regions defined by hills and/or river valleys (**Figure 2a**). This structure is not an artifact of the denser sampling in this region, as it was still evident in an analysis after sub-sampling (Methods; **Supplementary Figure 2**). Highly localised structure is also seen in other parts of Spain, including four clusters within the Basque regions (**Figure 2b**), and a cluster that is exclusive to a ∼50 Km segment of the River Ebro in La Rioja (**Figure 1c**).

**Figure 2.**
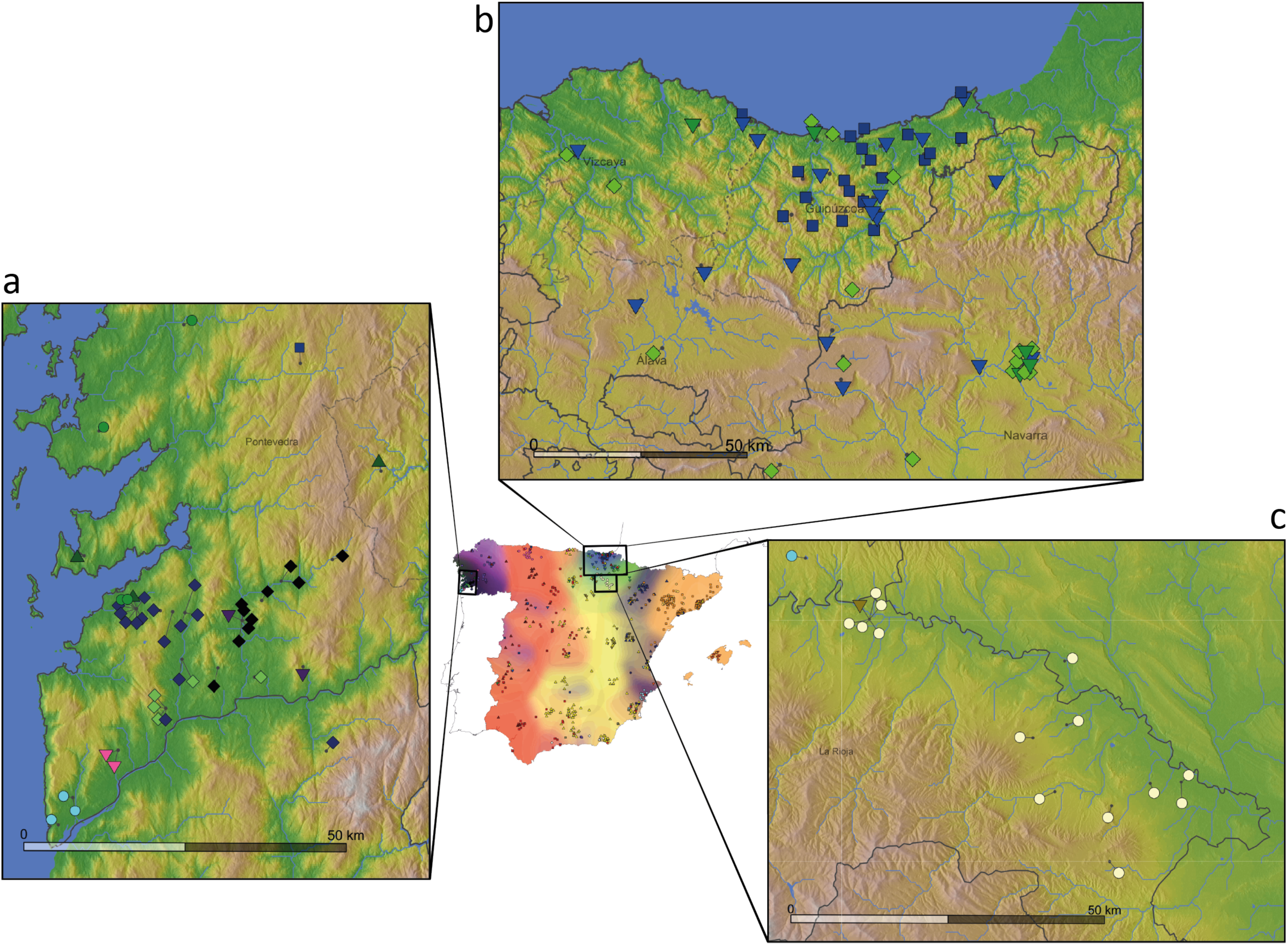
Ultra-fine-scale genetic structure within Spain. Points representing individuals are placed on each of the magnified maps and coloured as described in Figure 1, with short dark lines pointing to their precise locations (the average birthplace of their grandparents). The three magnified maps show local elevation, rivers, and water bodies, as well as borders of Autonomous Communities (solid black lines) and provinces (dashed lines and text). **(a)** Locations of individuals within the genetic clusters centred in Galicia. Note that we show this region at a higher level of the tree (14) as the lower level yields clusters with fewer than 3 individuals with fine-scale geographic location data. **(b)** Individuals within the clusters centred in the Basque-speaking regions of Pais Vasco (Basque Country) and Navarre. For visual clarity we only show the individuals that are within the clade coloured blue and green in Figure 1. This clade makes up the majority of all individuals located in this region, and a majority of this clade is located in this region. **(c)** Locations of individuals in a cluster exclusive to a ∼50km-wide region along the banks of the River Ebro in La Rioja (just south of Pais Vasco and Navarra).

To further understand the relationships between the clusters inferred by fineSTRUCTURE, we examined patterns in the matrix of ancestry sharing (‘coancestry’) between each pair of 1,413 individuals (**Supplementary Figure 3a**). In general, coancestry between individuals within a cluster is higher than between individuals in different clusters, reflecting genetic drift unique to each cluster. This effect is strongest for highly localised clusters, such as those in Galicia and País Vasco and La Rioja (**Supplementary Figure 3b**). These clusters also tend to have greater certainty in their cluster assignment (**Supplementary Figure 8b**). In contrast, the cluster labelled ‘central’ (shown with yellow triangles in **Figure 1b**) shows no clear drift signal. In fact, individuals in this cluster have – on average – more coancestry with the members of Basque-centred clusters (blue squares and triangles) than they do with other individuals in their own cluster (*p* < 0.02; **Supplementary Figure 3c**). Theoretical arguments predict (Methods) that this effect can only occur if admixture from a highly drifted group into another population takes place, and implies directionality of this admixture. Thus, this signal provides evidence of admixture into the ‘central’ cluster from a group related to the Basque populations.

Next, we sought to characterize the relationship between Iberians (combining Spanish and Portuguese individuals) and non-Iberian groups, to understand the extent to which recent migrations from outside Iberia have influenced modern-day DNA in Spain. We constructed a combined dataset (300,895 SNPs) of 2,920 individuals from Spain, Europe, north Africa^18^ and sub-Saharan Africa^19^ (Methods). We used fineSTRUCTURE to identify 29 non-Iberian ‘donor groups’ (Methods; **Supplementary Figure 4a**). We extended the fineSTRUCTURE model to re-cluster individuals within Iberia, now based only on their levels of ancestry sharing across these 29 groups (Methods). These clusters capture the impact of migration into and across Spain, removing the effects of simple isolation events.

Using this approach we inferred six distinct clusters within Iberia (**Figure 3a**), many fewer than in the Spain-only analysis (**Figure 1a**), implying that much of the fine-scale structure seen within Spain is a result of regional genetic isolation. The six clusters still associate with geographical regions, predominantly in the east-west direction rather than north-south. Notably, the extensive sub-structure in Pontevedra disappears, and indeed these individuals now co-cluster with Portuguese individuals. Therefore, the extensive fine-scale structure in Galicia is explained by local drift effects. In contrast, a distinct cluster still occurs within the Basque region. This indicates that alongside regional isolation, distinctive levels of ancestry sharing with non-Spanish groups contribute to fine-scale structure in this region.

**Figure 3.**
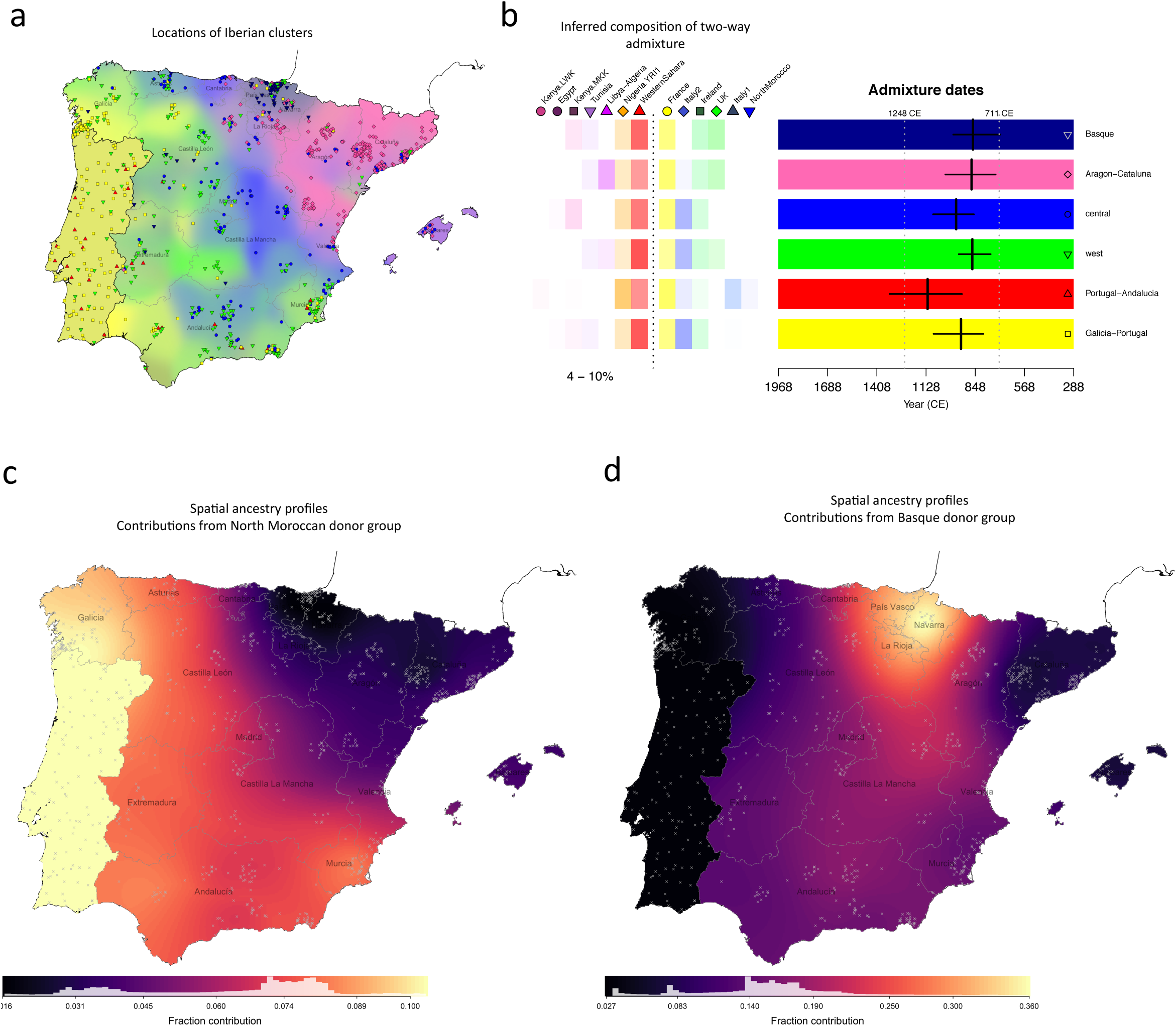
Characterising genetic contributions to Iberia. **(a)** Geographic distribution of Iberian individuals grouped into 6 clusters based on haplotype sharing with external populations (Methods). Background colours and the positions of points on the map are determined using the same procedure as for **Figure 1b**, with the exception of individuals of Portuguese origin. No fine scale geographic information was available for these individuals, so we placed them randomly within the boundaries of Portugal, and show a single background colour (Methods). **(b)** Admixture dates and mix of admixing groups in single-date, two-way admixture events, as inferred using GLOBETROTTER. On the left are the donor groups inferred to best represent the two ancestral populations involved in the admixture event (separated by a dashed line), along with the range of inferred admixture proportions of the smaller side. Estimated dates, and 95% bootstrap intervals are shown on the right, for each target group. Target groups are the set of Iberian clusters, as shown in (a). The white vertical dashed lines show the time of the initial Muslim invasion (711 CE) and the Siege of Seville (1248 CE), between which around half (or more) of Iberia was under Muslim rule. The admixture dates assume a 28-year generation time, and a ‘now’ date of 1940 (the approximate average birth-year of this cohort). Detailed results of this GLOBETROTTER analysis are tabulated in Supplementary Table S3a. **(c) and (d)** We estimated ancestry profiles for each point on a fine spatial grid across Spain (Methods). Gray crosses show the locations of sampled individuals used in the estimation. Map (c) shows the fraction contributed from the donor group ‘NorthMorocco’. Map (b) shows the fraction contributed from the donor group ‘Basque1’, which we defined based on the Spain-only fineSTRUCTURE analysis (**Figure 1a**). Maps for other donor groups are shown in **Supplementary Figure 5**.

To characterise the genetic make-up of these six Iberian clusters we estimated their ‘ancestry profiles’. We fitted each cluster as a mixture of (potentially) all 29 donor groups to approximate the unknown ancestral groups that actually contributed to modern-day Iberian individuals (Methods). This approach accounts for the stochasticity in ancestral relationships along the genome and was previously shown to be informative in the context of the British Isles^6^. Only six of the 29 donor groups show a contribution > 1% in Iberia, and all are located in Western and Southern Europe, and north-west Africa (**Supplementary Figure 4**). For all six Iberian clusters the largest contribution comes from France (63 - 91%), with smaller contributions that relate to present-day Italian (5 - 17%) and Irish (2 - 5%) groups. With the exception of the Basque cluster, these three donor groups dominate, and contribute proportionally similar amounts throughout Iberia, so probably represent ancient ancestry components rather than recent migration. In contrast, north Moroccan ancestry shows strong regional variation (**Figure 3c**, Methods). See Supplementary Information for a fuller discussion of the ancestry profiles.

To distinguish between possible scenarios that could produce these patterns, we applied the GLOBETROTTER method^20^ to each of our six clusters (Methods). GLOBETROTTER infers dates of admixture and the make-up of the source populations, and tests whether admixture patterns are consistent with a simple mixing of two groups at a single time in the past, compared to more complex alternative models. GLOBETROTTER found strong evidence (*p* < 0.01) of admixture for all six clusters (Methods; Table S3a in Supplementary Information). For all six clusters, an extremely similar event was inferred (**Figure 3b**), in a tight time-range of 860 - 1120 CE, and with similar source groups, present in varying proportions (4 - 10% for the minor group). The major source was inferred to contain almost exclusively European donor groups, and the minor source is made up of mainly north west African donor groups, including Western Sahara, and to a lesser extent west Africans (YRI), consistent with the overall ancestry profiles. The ‘Portugal-Andalucia’ cluster shows the greatest YRI contribution, and also shows some evidence of a second admixture date, with a more recent event involving only sub-Saharan-African-like and European-like source groups (see Supplementary Information for a fuller discussion; **Supplementary Figure 6**). This indicates a recent pulse of sub-Saharan African DNA, independent of the north African component.

GLOBETROTTER shows a subtle preference for Western Sahara as a source of North African DNA, as opposed to north Morocco. This might be explained if modern-day north Moroccan haplotypes are more similar to present-day Spanish individuals than the historical admixing population was. Indeed, previous work^18^ found that modern-day north Moroccan haplotypes carry a component of European ancestry that most likely arrived subsequent to the detected admixture event, and this is consistent with a mixture analysis we performed of the north Moroccan group itself (**Supplementary Figure 7**; Methods).

Our earlier results imply the incorporation of Basque-like DNA elsewhere within Spain. We next incorporated the clade labelled ‘Basque1’ as a potential donor group, to characterise and date this event (Methods). Ancestry profiles show contributions of Basque-like DNA (**Figure 3c**) highest in places immediately surrounding the main location of the Basque donor group (País Vasco), and much higher southwards than to the east and west. GLOBETROTTER yields congruent results, and inferred dates for the arrival of Basque-like DNA in the range 1190 – 1514 CE, more recent than the north west African influx (**Supplementary Figure 10**).

## 3 Discussion

Our observation that genetic differences are small in the north-south direction within Spain, and evidence of gene flow preferentially in this direction, are most straightforward to interpret in the light of historical information regarding the *Reconquista,* during which Christian-controlled territory in the north moved gradually southwards from the mid-8^th^ Century, following the Muslim invasion of Iberia (711CE). By 1249 almost all of Iberia was under Christian rule, and the Battle of Granada in 1492 marks the end of Muslim rule in Iberia. There is historical evidence of migration of peoples from the northern Christian kingdoms into newly conquered regions during the *Reconquista*^10,21^. The east-west boundaries of the clusters we see in the north of Spain correspond closely to the regions of broad linguistic differences in the Christian-ruled north, which date back to at least the first 200 years of Muslim rule (Figure 1c), and we date the southwards movement of Basque DNA later within the *Reconquista* itself. Thus, it appears that present-day population structure within Spain is shaped by population movements within this key period.

We also detect a genetic footprint of the Muslim invasion, and subsequent centuries of Muslim rule, itself. Following the arrival of an estimated 30,000 combatants^22^, a civilian migration of unknown numbers of people occurred, thought to be mainly Berbers from north Morocco and settling in many parts of the peninsula^22^. Our analysis, alongside previous smaller studies^12,23^, imply a substantial and regionally varying genetic impact. Our results further imply that north west African-like DNA predominated in the migration. Moreover, admixture mainly, and perhaps almost exclusively, occurred within the earlier half of the period of Muslim rule (**Figure 3b**). Within Spain, north African ancestry occurs in all groups, although levels are low in the Basque region and in a region corresponding closely to the 14^th^-century ‘Crown of Aragon’ (**Figure 3c**). Therefore, although genetically distinct^16,22,23^, this implies that the Basques have not been completely isolated from the rest of Spain over the past 1300 years.

Perhaps surprisingly, north African ancestry does not reflect proximity to north Africa, or even regions under more extended Muslim control. The highest amounts of north African ancestry found within Iberia are in the west (11%) including in Galicia, despite the fact that the region of Galicia as it is defined today (north of the Miño river), was never under Muslim rule^24^ and Berber settlements north of the Douro river were abandoned by 741. This observation is consistent with previous work using Y-chromosome data^12^. We speculate that the pattern we see is driven by later internal migratory flows, such as between Portugal and Galicia, and this would also explain why Galicia and Portugal show indistinguishable ancestry sharing with non-Spanish groups more generally. Alternatively, it might be that these patterns reflect regional differences in patterns of settlement and integration with local peoples of north African immigrants themselves, or varying extents of the large-scale expulsion of Muslim people, which occurred post-*Reconquista* and especially in towns and cities^10,21^.

We show that population structure exists at ultra-fine scales in Galicia (**Figure 2a**), particularly in the province of Pontevedra, with some clusters having geographic ranges of less than 10 Km. To our knowledge, these results represent the finest scales over which such structure has yet been observed in humans. It will be interesting to identify whether (and if so which) other parts of the world show similar patterns. Pairs of individuals within these clusters show high levels of coancestry relative to the rest of Spain (**Supplementary Figure 3b**). In contrast, when we only consider their patterns of coancestry with non-Iberian groups this structure disappears: individuals from Pontevedra are indistinguishable from those from Portugal and other parts of western Spain (**Figure 3a**). Therefore, the very strong population stratification observed in Galicia can most easily be explained through very recent geographic isolation, occurring subsequent to major migrations into the region (see Supplementary Information for further discussion).

Overall, the pattern of genetic differentiation we observe in Spain reflects the linguistic and geopolitical boundaries present around the end of the time of Muslim rule in Spain, suggesting this period has had a significant and long-term impact on the genetic structure observed in modern Spain, over 500 years later. In the case of the UK, similar geopolitical correspondence was seen, but to a different period in the past (around 600 CE)^6^. Noticeably, in these two cases, country-specific historical events rather than geographic barriers seem to drive overall patterns of population structure. The observation that fine-scale structure evolves at different rates in different places could be explained if observed patterns tend to reflect those at the ends of periods of significant past upheaval, such as the end of Muslim rule in Spain, and the end of the Anglo-Saxon and Danish Viking invasions in the UK.

It appears that within-country population structure occurs across the world^6,25^, and in this study is observed down to scales of less than 10km. Such strong, localised genetic drift predicts the existence of geographically localised rare mutations, including pathogenic ones. For example, cases of a specific form of inherited ataxia (SCA36) cluster within a specific region of Galicia^26^. Our results imply this type of phenomenon will tend to happen in specific areas, such as Galicia in the case of Spain, and can arise even where population density is high and in the absence of obvious strong geographic barriers.

## 5 Acknowledgements

We acknowledge support from the Wellcome Trust (203141/Z/16/Z, 090532/Z/09/Z, 098387/Z/12/Z, 095552/Z/11/Z) and Fondo de Investigación Sanitaria (Grants PI13/01136 and PI16/01057). We thank G. Hellenthal, D. Lawson, and G. Busby for advice on the use of fineSTRUCTURE and GLOBETROTTER. Also G. Hellenthal for providing computer code for the analysis of cluster assignment uncertainty in the fineSTRUCTURE analysis. We also thank F. Dubert García and R. Villares Paz for advice on relevant historical sources. The support of the Spanish National Gentyping Center (CEGEN-PRB2) is also acknowledged with appreciation.

## 6 Author contributions

A.C., P.D., and S.M. initiated and designed the study. C.F-R., C.R-P., and I.Q-G. collected the data for the Spanish cohort and carried out genotype calling under the direction of A.C.. C.B. performed the analyses under the joint supervision of S.M. and P.D. A.C. provided historical information. C.B., A.C, P.D., and S.M. wrote the manuscript.

## 7 Ethics statement

The collection of the Spanish genotype data was approved by the “Comité Ético de Investigación Clínica de Galicia”, and each of the institutional review boards of the participating hospitals. All samples were obtained with written informed consent reviewed by the ethical board of the corresponding hospital, in accordance with the tenets of the Declaration of Helsinki. A previous analysis of these data was published in 2013^27^. All other data used in the paper were previously published and are publicly available for research use.

## 9 Methods

### Data and quality control

For the Spain-only analysis we used genotype data that was originally collected and typed for a colorectal cancer GWAS^27^. Biological samples were sourced from a variety of hospitals across Spain as well as the Spanish National DNA Bank. All samples were assayed together by Affymetrix^1^ in the same facility. See^27^ for full details of sample collection and genotype calling. We used both cases and controls, which totaled 1,548 individuals prior to quality control. Individuals from all 17 of Spain’s Autonomous Communities are represented in this dataset, but the Spanish territories of Melilla, Ceuta, and the Canary Islands are excluded from analyses involving geographic labelling due to limited sampling in these areas (4 samples).

After applying a series of quality control filters (Supplementary Information) we phased the genotype data using SHAPEIT v2^28^ with a reference panel and recombination map from Phase I 1000 Genomes^29^. Close relatives (kinship coefficient > 0.1), and individuals with evidence of recent, non-Spanish ancestry were included in the phasing, but excluded from the fineSTRUCTURE analysis. This procedure resulted in 1,413 samples and 693,092 SNPs for our main analysis.

For all analyses involving individuals from outside of Spain we combined four sources of genotype data: European samples from POPRES^30^, north African samples from^18^, sub-Saharan African samples from Hapmap Phase 3^19^, and the Spanish samples described above. The POPRES and north African samples were typed on the Affymetrix 500K array, which overlaps substantially with the Affymetrix 6.0 array. We first merged these data and then applied quality filters to the combined dataset of 6,617 individuals, which we then phased altogether. After excluding related individuals, those with self-reported mixed ancestry, and sub-sampling some heavily sampled regions (e.g. Switzerland) the data set comprised 2,920 individuals and 300,895 SNPs, which we used in our joint analyses. See Supplementary Information for full details of quality control and phasing.

### Clustering using fineSTRUCTURE

We inferred clusters of individuals based on genetic data only by applying the fineSTRUCTURE method^17^, which uses a model-based approach to cluster individuals with similar patterns of shared ancestry. Within the fineSTRUCTURE framework, shared ancestry is measured as the total amount of the genome (in centiMorgans (cM)) for which individual *i* shares a common ancestor with individual *j*, more recently than all the other individuals in the sample. This is estimated for each pair of individuals *i* and *j*, defining a square matrix referred to as the ‘coancestry’ matrix (e.g. **Supplementary Figure 3a**). This matrix is then used to cluster individuals into groups with similar patterns of coancestry, i.e. similar rows and columns in the matrix. We applied fineSTRUCTURE using the procedure recommended by the authors, except in one aspect: to measure shared ancestry (coancestry) we used the total amount of genome (in cM) for which individual *i* shares a common ancestor with individual *j*, more recently than all the other individuals in the sample. The software default coancestry measure is the number of contiguous segments (‘chunks’) rather than the total amount of genome, but we found the alternative measure to be more robust to artefacts such as genotyping error (**Supplementary Figure 9**; Supplementary Information).

Using the coancestry matrix, fineSTRUCTURE applies a Markov chain Monte Carlo (MCMC) procedure to find a high posterior probability partition of individuals into a set of clusters. The number of clusters is not specified in advance, but rather estimated under the fineSTRUCTURE probability model. Having found a set of clusters, fineSTRUCTURE then infers a hierarchical tree by successively merging pairs of clusters whose merging gives the smallest decrease to the posterior probability (of the merged partition) among all possible pairwise merges.

We ran several fineSTRUCTURE analyses using phased haplotypes from the following sets of individuals:

(A) Spanish individuals
(B) Spanish and Portuguese individuals
(C) Non-Spanish individuals

In all cases we used CHROMOPAINTER software (v2)^17^ to estimate the coancestry matrix, followed by fineSTRUCTURE’s clustering and tree-building procudures. We used a previous successful application of fineSTRUCTURE^6^ as a guide for the number of iterations in the MCMC, and other required parameters (see Supplementary Information for full details). We also checked that the MCMC samples were largely independent of the algorithm’s initial position by visually comparing the results of two independent runs starting from different random seeds. Good correspondence in the pairwise coincidence matrices of the two runs indicates convergence of the MCMC samples to the posterior distribution^17^. See, for example, **Supplementary Figure 8a** showing the two independent runs for analysis (A). Without loss of generality, we used the first of these two runs in our main analysis.

### The statistical uncertainty of cluster assignments

For analysis (A) we measured uncertainty in the assignment of individuals to clusters by using a procedure described formally in^6^, which uses the information from multiple samples of the fineSTRUCTURE MCMC. Informally, the procedure measures the overlap between a cluster *k*, and individual *i*’s assigned cluster in each of the MCMC samples within a single fineSTRUCTURE run. This can take values between 0 and 1, and sums to 1 across all clusters for a given individual. It provides a measure of certainty about the assignment of individual *i* to each cluster *k* in the final set of clusters. The ‘cluster assignment certainty’ for an individual is the value corresponding to the final cluster assignment, and will be close to 1 if they are assigned to a cluster with largely the same set of individuals in each MCMC sample. This measure can be obtained for different layers of the hierarchical tree by summing the values for the clusters that merge at each layer.

For fineSTRUCTURE analysis (A), cluster assignments at higher levels of the hierarchical tree are typically more certain than lower-level clusters. At the broader level of the tree shown (14 background colours in **Figure 1b**) 94% of individuals have a cluster assignment certainty greater than 0.7; and at the finer level (shown with points in **Figure 1b**), this level of certainty is reached by 89% of individuals (see **Supplementary Figure 8b**). Furthermore, the clusters with the highest certainty tend to be those with greater geographic localization, e.g. those labelled ‘LaRioja’ and ‘Baleares’ (**Figure 1**).

### Selection of levels of the Spanish tree to analyse

In the fineSTRUCTURE analysis (A) 145 clusters were inferred. In order to examine properties of these clusters, such as their geographical locations, we focused on two different levels of the tree, thus highlighting both broad-scale and fine-scale structure. There is no ‘right’ level of the tree to pick, but we chose them based on the sizes of the clusters. To examine broad-scale structure we chose a level (14 clusters) such that all clusters were larger than size 20. Recall that moving up the tree, at each level a pair of clusters is merged. At the base of the tree, newly separated groups typically contain few (<15) individuals, with higher clustering uncertainty. To avoid the presence of such minor clusters, we traversed up the tree until the first time a merge occurred between two larger clusters (>15 individuals). This occurs at the level of 49 clusters. However, at this level over half (28) of the clusters are within the clade involving individuals mostly from south-west Galicia (labelled ‘Galicia_Pontevedra’ in **Figure 1a**), and for many of these clusters fine-scale geographic information was only available for one or two individuals, and/or the cluster contained fewer than 15 individuals. Therefore, to aid visualization in **Figure 1b** and **Figure 2a** we only show the clusters at the higher level of the tree (level 14) for *this* clade, although they are still shown in **Figure 1a**.

### Map-based data visualisation

Geographic information was available for individuals in the Spanish cohort, along with their age at collection, sex, and genotyping plate (controls only) and batch used in genotype calling. The geographic information includes region of origin (Autonomous Community) for all individuals, and for 953 individuals (65%) the birthplace (town) of all four grandparents. We assigned each individual to a geographic coordinate by matching the text (e.g ‘Barcelona’) to a municipal region as defined by the Spanish Statistical Office (**www.ine.es,** 2014) and coding them to the centre of the matching region. Some locations were not themselves a municipality, so we coded these individuals to the centre of the nearest municipality, identified by using Google Maps. However, in the fineSTRUCTURE analysis we clustered all individuals, i.e. also included those individuals for whom the exact birthplaces of their grandparents was not known, but these are not used in interpreting the spatial distribution of the inferred genetic structure.

In figures showing a map of Spain (e.g. **Figure 1b**) each individual is represented by a point placed at the average coordinate (centroid) of their grandparents’ birthplaces (coordinates were derived as described above). Where many individuals have the same coordinate, such as in Barcelona, points have been randomly shifted, by no more than 24 Km, to aid visualisation. To visually represent the discrete assignment of individuals to clusters by fineSTRUCTURE, the members of each cluster are represented using the same colour and symbol. We also coloured the background of the maps using a Gaussian kernel smoothing procedure on a regular grid of 3 Km-wide squares across Spain. Informally, each square in the grid is coloured according to the relative contributions of each cluster, where contributions are measured by Gaussian densities centred on the location of each individual. See Supplementary Information for a formal description.

### Signals of drift and admixture in the coancestry matrix

Recall that coancestry (as we have used it) measures the amount of genome (in cM) for which an individual *i* shares its most recent common ancestor with another individual *j*, out of all the individuals in the sample. Properties of this matrix are informative of patterns of drift and admixture within and across clusters inferred by fineSTRUCTURE. Specifically, excess coancestry between individuals in the same cluster (within-cluster coancestry) is a natural measure of genetic drift of that cluster relative to all the other clusters^17^. In general, individuals are often observed to have the highest levels of coancestry with other individuals in their assigned cluster. This is not a constraint of the fineSTRUCTURE model; rather it is because if two individuals have similar patterns of shared ancestry, they are naturally also likely have more recent shared ancestry between them. However, it is possible for this not to be the case, and this is informative of admixture (see Supplementary Information for further discussion).

We looked for such signals in the case of Spain by testing whether a cluster (inferred by fineSTRUCTURE) has a within-cluster coancestry that is, on average, smaller than its coancestry with another cluster. We used the 26 clusters (indicated with symbols on the axes in **Supplementary Figure 3a**), which contained at least 13 individuals. To avoid potential bias due to uneven sizes of the clusters, we estimated within-cluster coancestry levels by randomly sub-sampling (without replacement, as coancestry is only defined between two different individuals) each of the 26 clusters such that there were of equal size (13). We re-computed the coancestry matrix using CHROMOPAINTER, and the same set of parameters as in fineSTRUCTURE analysis (A), but using this smaller subset. We repeated this 200 times and used these resamples to compare coancestry levels across clusters. For each resample and each cluster we computed the mean of the coancestry values within that cluster (excluding zeros on the diagonals), and with each of the other clusters. We then computed a *p*-value using the number of resamples (*S*) for which the mean within-cluster coancestry is smaller than the mean coancestry with each of the other clusters. That is: 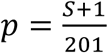 Results are shown in **Supplementary Figure 3b** and **c**.

### Defining ‘donor groups’

In order to define a set of ‘donor groups’ using the combined European, north African, and sub-Saharan African individuals, we applied fineSTRUCTURE as described above in several rounds, each using the following sets of individuals:

(CI) All individuals combined (excluding Spanish but including Portuguese)
(CII) Individuals from north Africa only
(CIII) Individuals from Europe only.

In analysis (CI) fineSTRUCTURE cleanly split the 3 main groups corresponding to Europe, north Africa, and sub-Saharan Africa, as well as inferring finer sub-structure. However, in order to maximise the power to detect finer scale structure^17^, we obtained fineSTRUCTURE results for the north African and European groups independently. That is, using coancestry matrices that only allow copying within each set of individuals in (CII) and (CIII), respectively. We then defined a set of donor groups based on the clusters and hierarchical trees inferred by fineSTRUCTURE in these three analyses. We considered the following factors in defining donor groups. Ideally, each donor group would contain about the same number of samples, and not be too small. Donor groups should also be relatively homogeneous with respect to their shared ancestry with the population of interest (in this case Iberia), although could be heterogeneous within themselves. We therefore prioritised donor group size over capturing finer scale structure that might exist within donor groups themselves. See Supplementary Information for full details. Our procedure resulted in a total of 29 donor groups with median size 30, and minimum size 16, and their locations are shown in **Supplementary Figure 4a**. Labels of the inferred groups are based on the sampling locations of most of the individuals in a given group. In some cases the majority of individuals were split across two locations, and this is indicated by a multi-region label (e.g. Germany-Hungary).

### Treatment of Portugal

One cluster in the fineSTRUCTURE analysis (CIII) overlaps significantly (98%) with the individuals with grandparental origins in Portugal as reported by the data source (POPRES). For the purposes of the analyses in this chapter (and fineSTRUCTURE analysis (B)), this group of 117 individuals is referred to as ‘Portugal’ or ‘Portuguese individuals’ (e.g. in **Supplementary Figure 4a**). The strong genetic similarity between individuals from Portugal and Spanish individuals, especially those located in Galicia (**Supplementary Figure 1a**), means they are likely to share a similar admixture history, and including Portugal as a *donor* group would mask the signal from those shared events. We therefore excluded them from the set of donor groups and instead treated them in the same way as the Spanish individuals. This is analogous to the rationale for excluding Ireland as a donor group in the British Isles study ^6^.

### Clustering on the basis of haplotype sharing with external groups

Here we describe the method we used to infer clusters of Iberian individuals with distinct patterns of haplotype sharing with external groups. We used the fineSTRUCTURE clustering algorithm but with a modified version of the input coancestry matrix. Specifically, we compute the coancestry (using CHROMOPAINTER) between each Iberian individual and each of the non-Iberian individuals, as described above, but only allowing Iberian individuals to copy from non-Iberian individuals. This results in a rectangular matrix, *X*, of size *N × M*, where *N* is the number of Iberian individuals and *M* the number of non-Iberian individuals. We then constructed an (*N*+*M*) *x* (*N*+*M*) square matrix *C*, such that,

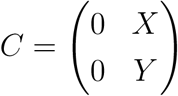

and matrix *Y* contains zeros, except for each of the (block) diagonal entries corresponding to pairs of individuals within the same donor population *k*. These entries each take value the *gk*, which is determined such that the mean of all the entries corresponding to donor population k are the same for the sub-matrix *X* as the sub-matrix *Y*. The zeros in the matrix *C* have the effect of not allowing any copying from or to the Iberian individuals to contribute to the fineSTRUCTURE likelihood. We then run fineSTRUCTURE algorithm using the ‘force file’ option (-F), where each ‘continental’ group is a donor group, thus only allowing splits and merges to take place among Iberian individuals. We used a *c*-factor of 0.0579, which was computed in the manner described in Supplementary Information, but using segments of DNA from a CHROMOPAINTER run where we only allowed Iberian individuals to copy from non-Iberian individuals.

### Estimating ancestry profiles

We estimated ancestry profiles for each of the Iberian clusters using the procedure described in ^6^. Briefly, we use CHROMOPAINTER to compute a coancestry vector for each Iberian individual, where we only allow them to copy from haplotypes in the donor groups (as with matrix *X* above), and then average the coancestry vectors within each cluster and donor group. For each Iberian cluster *i*, we then find a smaller set of donor groups which together (as a non-negative linear mixture whose coefficients sum to 1) best explains its cluster-averaged coancestry vector, 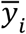.(see Supplementary Information for full details). The vector of coefficients in the linear mixture is the ancestry profile for cluster *i*, and its elements sum to 1. Results for six Iberian clusters are shown in **Supplementary Figure 4b**. We also performed a complementary analysis where we treated each *donor group* in turn as.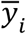., after removing their corresponding elements in the coancestry vectors and re-normalising (**Supplementary Figure 7**; Supplementary Information).

We measured uncertainty in these ancestry profiles by re-estimating the cluster-averaged coancestry vectors using a set of pseudo individuals. Each pseudo individual is formed by randomly selecting an individual in cluster *i*.for each chromosome, and summing the observed chromosome-level coancestry vectors across all chromosomes. We then compute 1000 such re-estimations and report the range of the inner 95% of the resulting bootstrap distribution.

### Spatially smoothed ancestry profiles

The availability of fine-scale geographic information for many of the Spanish individuals allowed us to estimate the spatial distribution of shared ancestry (**Figure 3c** and **d**). Instead of averaging coancestry over individuals within a cluster, we average across geographic space using a Gaussian kernel smoothing method that varies the kernel band-width depending on the density of available data points (see Supplementary Information for details). This gives a set of coancestry vectors, 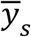, for each grid-point s in a fine spatial grid across Spain. We then compute ancestry profiles for each of the grid-points in the same way as for the Iberian clusters (described above), but setting 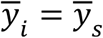instead of a cluster-averaged coancestry vector. We visualize the results by colouring each grid point according to the value of its coefficient for a single donor population of interest (e.g. NorthMorocco in **Figure 3c**). For any grid-point s. and individual *i*.located within the borders of Portugal we set all 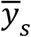to be the average coancestry vector across these individuals, because we have no fine-scale geographic information for them. This means in Portugal there is always one colour plotted.

### Estimating admixture dates and source populations

We used the GLOBETROTTER algorithm to estimate dates, proportions and configurations of admixture events^20^. Briefly, GLOBETROTTER uses ‘paintings’ from the CHROMOPAINTER algorithm to construct ‘coancestry curves’. These curves measure the rate of decay of linkage disequilibrium with genomic distance, between sites with ancestry from a pair of source populations. The parameters of exponential functions fitted to these curves (decay rates and intercepts) are used to estimate admixture dates, admixture proportions, and the best fitting mix of modern-day groups that characterise the ancestral populations involved in an admixture event.

We conducted two analyses using GLOBETROTTER. The first (gtA) was designed to detect admixture event(s) in the history of Iberia that might involve any combination of non-Iberian source populations, without any prior assumptions on the nature of the event. The second analysis (gtB) was designed to detect only admixture event(s) involving a Basque-like source population, i.e. based on a prior hypothesis. In each case we defined a set of target groups within which to look for an admixture event; a set of donor groups, which we allow to be donors in the initial ‘painting’; and a set of surrogate populations, which we allowed GLOBETROTTER to consider as components of any admixture event (Supplementary Information Table S2).

After identifying the presence of admixture based on criteria recommended by the authors, we next evaluated the evidence for more complex admixture events (e.g. multiple dates or more than two source populations). GLOBETROTTER automatically tests for these using a series of criteria based on how well the coancestry curves fit the models for different types of admixture scenarios^20^. Using GLOBETROTTER’s automated criteria, in analysis (gtB) there was no evidence that a two-date admixture model fitted better than a one-date model. However, for some target populations in analysis (gtA) there was some evidence for a two-date admixture event, although the one-date event fit well in all cases (Supplementary Information Table S3a). Given the potentially complex nature of admixture in Iberia, we further evaluated the evidence for a two-date admixture event by considering the model fits for each coancestry curve separately (**Supplementary Figure 6**; Supplementary Information). Notably, only the coancestry curves involving a sub-Saharan African surrogate group fit better to a two-date admixture event. The improved fit for the curve for the sub-Saharan African surrogate group ‘Nigeria.YRI1’ is visually apparent in the coancestry curve shown in **Supplementary Figure 6**. We therefore consider the one-date admixture event to be a better fit overall, but that there is some evidence for a second event involving sub-Saharan African-like DNA mixing with European-like DNA, with the strongest evidence for this in the Iberian cluster, ‘Portugal-Andalucia’. In the target groups where there is evidence of this, GLOBETROTTER infers dates in the range 1370 - 1700 CE (assuming a 28-year generation time).

## Footnotes

1 Now part of Thermo Fisher Scientific.

